# TET1 non-catalytic activity shapes the chromatin landscape associated with de novo methylation establishment in the male germline

**DOI:** 10.1101/2025.10.29.685418

**Authors:** R. D. Prasasya, J. J. Kim, Z. Liu, R. M. Kohli, M. S. Bartolomei

**Author notes:** Correspondence (MSB).

## Abstract

In the germline, DNA methylation is globally erased in primordial germ cells (PGCs), enabling establishment of sex-specific methylomes in prospermatogonia or oocytes. The catalytic activity of TET1 is required for complete demethylation in PGCs, yet sperm from *Tet1^−/−^* mice display methylation defects not explained by incomplete erasure. Instead, these defects arise from abnormal *de novo* methylation during development, coinciding with erosion of H3K4me3, a chromatin modification that blocks DNMT3A/3L. Using a catalytically inactive *Tet1^HxD^* mouse line, we demonstrate a non-catalytic role of TET1 in promoting H3K4me3 deposition and protecting a subset of sperm hypomethylated regions from aberrant methylation in prospermatogonia.

## Introduction

DNA methylation is one of the most sexually dimorphic epigenetic marks in mammalian gametes (Stewart et al. 2016). Following specification from cells of the proximal epiblast, primordial germ cells (PGCs) undergo epigenetic reprogramming that involves both chromatin remodeling and genome-wide DNA methylation erasure (Saitou et al. 2005; Smith and Meissner 2013; Saitou et al. 2011; Hajkova et al. 2002, 2008). In the mouse developmental timeline, the PGC genome attains a hypomethylated state by embryonic day (E)13.5 (Seisenberger et al. 2012). From this common basal epigenetic state, the sperm genome becomes highly methylated (∼90% overall methylation in the mouse) while the oocyte genome remains intermediate, with distinct hyper- and hypomethylated domains (Kobayashi et al. 2012; Hanna et al. 2018). In the male germline, acquisition of DNA methylation occurs in prospermatogonia between E15.5 and postnatal day 0. In contrast, *de novo* methylation occurs postnatally in oocytes when individual follicles are recruited for growth during each estrus cycle. Despite the distinct timing and the divergent resulting methylomes between the two sexes, *de novo* DNA methylation is facilitated by a common DNA methyltransferase (DNMT) complex: DNMT3A and its noncatalytic co-factor DNMT3L (DNMT3A/3L) (Kaneda et al. 2004; Kato et al. 2007; Bourc’his and Bestor 2004; Dura et al. 2022).

The mechanisms that target DNMT3A/3L to distinct loci in prospermatogonia versus the growing oocyte to generate sex-specific methylomes are still unclear. Biochemical studies suggest crosstalk between histone post-translational modifications (PTMs) and genome accessibility for DNMT3A/3L. For example, the PWWP domain of DNMT3A binds to di- or tri-methylated H3K36 (H3K36me2/me3) (Dhayalan et al. 2010). In the germline, H3K36me2 enriched domains are highly correlated with DNA methylation acquisition in prospermatogonia while transcription-coupled H3K36me3 deposition directs DNMT3A/3L and is the rate-determining chromatin environment in oocyte *de novo* methylation (Shirane et al. 2020; Chotalia et al. 2009; Stewart et al. 2015). In contrast, biochemical and *in vitro* studies have shown that methylation of H3K4 (H3K4me1/2/3) repels the ADD domain of DNMT3A and DNMT3L, inhibiting DNA methylation activity (Zhang et al. 2010; Ooi et al. 2007). In the male germline, DNA methylation acquisition occurs broadly, except at regions enriched for H3K4me2 at the prospermatogonia stage (Singh et al. 2013). Our previous data also confirmed the high congruency of H3K4me3 enrichment in prospermatogonia with sperm hypomethylated regions (HMRs, regions identified as methylation canyons using high coverage whole genome bisulfite sequencing of the sperm genome) (Prasasya et al. 2024). A functional study targeting writers for H3K4 methylation in the context of male *de novo* methylation has yet to be done, likely owing to the concern for potential functional redundancies of various histone methyltransferases expressed at high levels in prospermatogonia (Zhao et al. 2021). Additionally, how the prospermatogonia or the oocyte genome is patterned for H3K36me2/3 or H3K4me1/2/3 prior to gametic *de novo* methylation establishment remains unresolved.

In our previous investigation of the methylcytosine dioxygenase, Ten-eleven translocation 1 (TET1), in germline reprogramming, we described extensive methylation defects in *Tet1^−/−^* sperm beyond the previously described limited numbers of TET1-dependent loci (i.e. imprinting control regions -ICRs- and meiosis-associated gene promoters that require TET1 during PGC epigenetic reprogramming) (Prasasya et al. 2024; Yamaguchi et al. 2012, 2013; Hill et al. 2018; SanMiguel et al. 2018). TET enzymes iteratively oxidize 5-methylcytosine (5mC) to 5-hydroxymethylcytosine (5hmC), 5-formylcytosine (5fC), and 5-carboxylcytosine (5caC) (Ito et al. 2011). 5hmC is poorly recognized by maintenance DNMT1, while 5fC and 5caC can be excised by thymine DNA glycosylase (TDG) resulting in an abasic site, activating the base excision repair machinery to recover unmodified cytosine (Kohli and Zhang 2013). Using mouse models harboring 5hmC-stalling (diminished activity in generating 5fC/5caC) TET1^V^ and catalytically inactive TET1^HxD^, we established the necessity of TET1 iterative oxidative capability in germline epigenetic reprogramming. Intriguingly, only a small portion of methylation defects observed in *Tet1^−/−^* sperm can be attributed to the loss of TET1 catalytic activity. In fact, expression of full-length, catalytically compromised TET1 (TET1^HxD^) was sufficient to rescue about 2/3 of methylation defects observed in *Tet1^−/−^* sperm (Prasasya et al. 2024). In addition to the catalytic core located within the C-terminus, TET proteins possess an extensive N-terminus with the ability to interact with and recruit several histone modifiers such as OGT, Sin3a, and PRC2 *in vivo* (Vella et al. 2013; Zhu et al. 2018; Wu et al. 2011; Chrysanthou et al. 2022). Here, we demonstrate that the majority of hypermethylation in *Tet1^−/−^*sperm arises from abnormal *de novo* methylation and coincides with diminished H3K4me3 at intergenic hypomethylated regions. Importantly, catalytically inactive TET1 was sufficient to revert the majority of H3K4me3 defects observed in *Tet1^−/−^* germline to WT levels, establishing a non-catalytic role for TET1 in shaping the chromatin landscape of the male germline.

## Results and Discussion

We conducted global DNA methylation analysis of *Tet1^−/−^* and WT (*Tet1^+/+^*) PGCs at the completion of reprogramming, E13.5, using the mouse Illumina Infinium Methylation BeadChip. Consistent with previous reports, the *Tet1^−/−^*PGC genome was not globally hypermethylated compared to WT (Supplemental Fig. S1A) (Yamaguchi et al. 2013; Hill et al. 2018). Aberrantly methylated regions in *Tet1^−/−^* PGCs were almost exclusively hypermethylated and enriched for ICRs (Figure 1A, Supplemental Fig. S1B, Supplemental Table S1A; 1049 hypermethylated differentially methylated regions -DMRs-, 1 hypomethylated DMR; each DMR corresponds to a single probe on the array). Gene ontology (GO) analysis revealed that hypermethylated DMRs were enriched for genes involved in meiosis prophase I (Figure 1B), consistent with previous findings by us and others showing that promoter activation requires proper demethylation (Hill et al. 2018; Prasasya et al. 2024). We compared hypermethylated DMRs in *Tet1^−/−^* PGCs with differentially expressed genes (DEGs) from our previously described RNA-seq (Figure 1C, Supplemental Fig. S1C) (Prasasya et al. 2024). Only DMRs within promoters of meiosis-associated genes corresponded to downregulated expression (Supplemental Fig. S1C), suggesting that the transcriptional consequences of TET1-dependent hypermethylation outside of these meiosis-associated genes may manifest later in germ cell development.

**Figure 1.**
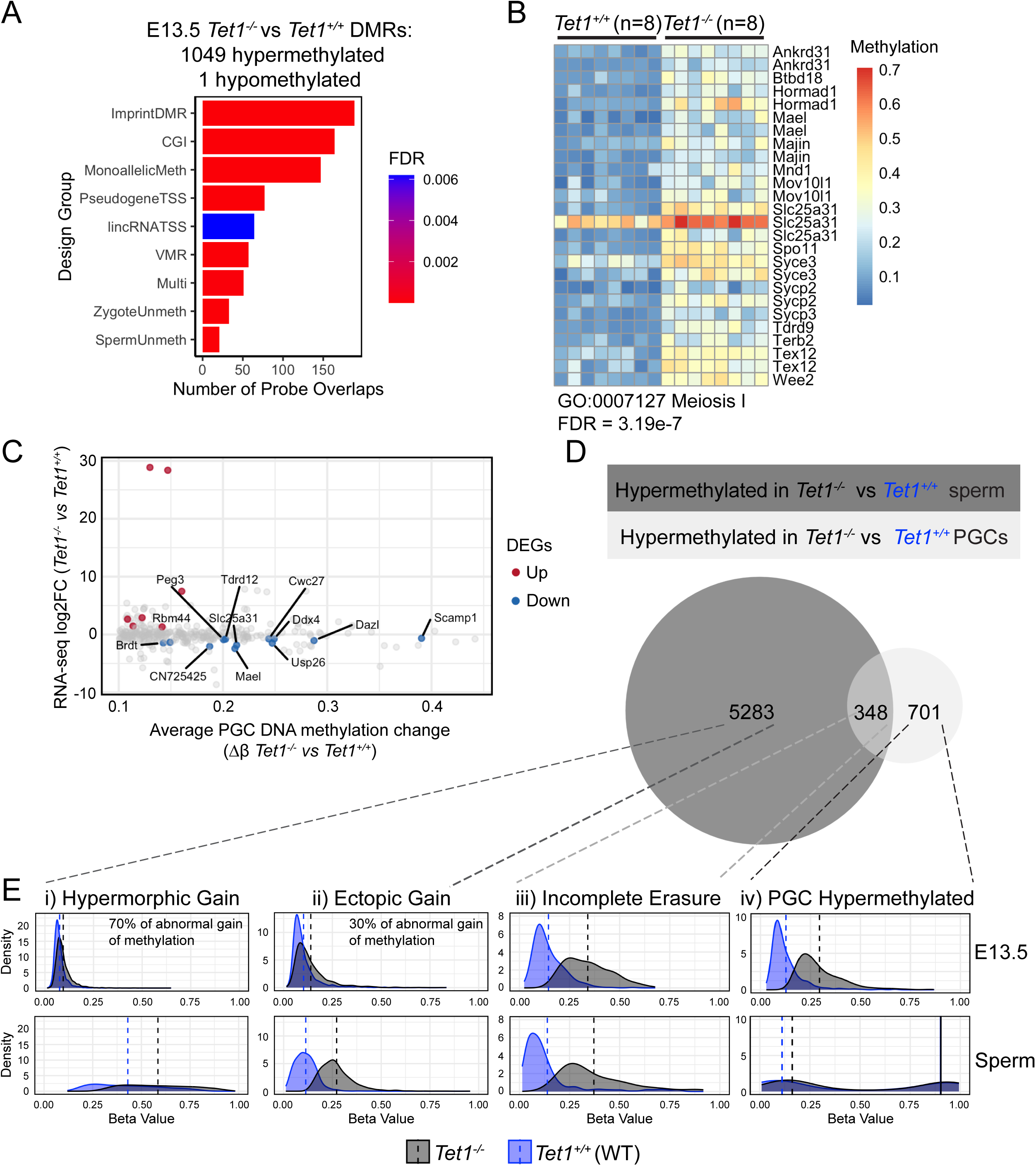
The majority of methylation defects in *Tet1^−/−^* sperm do not result from incomplete methylation erasure during epigenetic reprogramming. A) Top enriched design groups based on the Illumina Infinium Mouse Methylation BeadChip manifest for hypermethylated DMRs in *Tet1^−/−^* PGCs. B) Heatmap of probes associated with the most enriched GO term among hypermethylated DMRs in *Tet1^−/−^* PGCs. C) Scatterplot showing overlap between hypermethylated DMRs and DEGs in *Tet1^−/−^* PGCs. D) Comparison of hypermethylated probes (differentially methylated regions, DMRs: FDR ≤ 0.05; >10% methylation difference) in E13.5 *Tet1^−/−^* PGCs (n=8) vs *Tet1^+/+^* (WT, n=8) and those previously reported in *Tet1^−/−^*sperm (Prasasya et al. 2024). E) Methylation levels in E13.5 PGCs (top) or sperm (bottom) for *Tet1^−/−^* (black) and *Tet1^+/+^* (blue). DMRs are subsetted by origin of hypermethylation (i–iv). Average methylation levels are denoted by dotted vertical lines.

To assess whether incomplete DNA methylation erasure during PGC development explains the hypermethylation defects previously reported in *Tet1^−/−^* sperm (Prasasya et al. 2024), we compared hypermethylated DMRs at the two stages (Figure 1D-E, Supplemental Table S1B). 348 DMRs were hypermethylated in both *Tet1^−/−^* PGCs and sperm. As shown in the distribution plot (Figure 1E, panel iii, Supplemental Table S1E), average methylation levels of these regions were unchanged between sperm and PGCs for both genotypes, suggesting that only a small fraction (6.18%) of *Tet1^−/−^*sperm hypermethylation results from incomplete erasure in PGCs. In contrast, the majority of hypermethylated DMRs in *Tet1^−/−^* sperm showed no change in *Tet1^−/−^*PGCs compared to WT (Figure 1E, panel i-ii), indicating replication-coupled passive dilution of DNA methylation was sufficient to reprogram these loci. Instead, these regions abnormally acquired DNA methylation following successful erasure in *Tet1^−/−^* germ cells (Figure 1E, panel i-ii, Supplemental Table S1C-D).

When comparing WT PGCs and sperm, we noted a subset of regions where DMRs typically gain methylation during normal male germline development; in *Tet1^−/−^,* this acquisition was exaggerated, resembling a hypermorphic phenotype (Figure 1E, panel i). In contrast, a second subset of regions typically remained unmethylated in WT sperm but ectopically gained methylation in *Tet1^−/−^*(Figure 1E, panel ii). Out of the 5283 *Tet1^−/−^* sperm DMRs that originated from abnormal methylation acquisition, the majority belonged to the category of hypermorphic gain (3667 hypermorphic gain, 1602 ectopic gain, 14 sperm DMRs did not pass filter in the PGC data set). Finally, we identified 701 regions that required TET1 for reprogramming in PGCs but ultimately became fully methylated in both *Tet1^−/−^*and WT sperm (e.g. paternally methylated ICRs such as *H19* or *IG-DMR*) or sperm HMRs that continued to demethylate as the germ cells resumed mitosis in spermatogonial stem cells (SSCs) (Figure 1E, panel iv, Supplemental Table S1F). Design group enrichment analysis showed hypermorphic gain of DNA methylation following TET1 loss occurred most frequently at enhancers (Supplemental Fig. S1D), while ectopic gain or DNA methylation following TET1 loss most frequently occurred within CpG islands (Supplemental Fig. S1E). Regions with incomplete erasure occurred mainly at ICRs (Supplemental Fig. S1F, ImprintDMR) and those exclusively hypermethylated in *Tet1^−/−^* PGCs are most enriched for CpG islands (Supplemental Fig. S1G). Overall, these findings suggest that TET1 not only supports epigenetic reprogramming in PGCs but also regulates proper DNA methylation acquisition during male germline development.

We hypothesized that abnormal gain of methylation observed in the *Tet1^−/−^* male germline occurs concurrently with *de novo* methylation establishment and reflects changes in the chromatin landscape in prospermatogonia. To test this, we conducted CUT&RUN for H3K4me3, a histone PTM that inhibits DNMT3A/3L activity, in E17.5 prospermatogonia. We chose E17.5 as a mid-point of global wave of *de novo* methylation that occurs between E15.5 and birth in the male germline (Kobayashi et al. 2013; Seisenberger et al. 2012; Singh et al. 2013). For both WT and *Tet1^−/−^*, over 50% of identified peaks were located within distal intergenic regions (Supplemental Fig. S2A). Most identified peaks showed overlap between *Tet1^−/−^*and WT prospermatogonia, although *Tet1^−/−^* prospermatogonia had fewer consensus peaks compared to WT (67205 vs 95643 peaks; consensus peaks were defined as peaks called in at least 2 out of 3 biological replicates; Supplemental Fig. S2A, B, Supplemental Table S2A, B). Averaged signals across genic regions or intergenic peaks revealed distinct H3K4me3 changes in *Tet1^−/−^* prospermatogonia (Figure 2A). While *Tet1^−/−^* prospermatogonia showed higher overall H3K4me3 near transcriptional start sites (TSSs), H3K4me3 was markedly lower at intergenic peaks (Figure 2A). Differential enrichment (FDR ≤ 0.05, minimum 1.5-fold change vs WT) identified 22904 regions that lost H3K4me3, mostly intergenic regions, in *Tet1^−/−^*prospermatogonia (Supplemental Fig. S2C, Supplemental Table S3A). In contrast, 10971 regions that gained H3K4me3 mapped predominantly within genic regions (Supplemental Fig. S2C, Supplemental Table S3B). Notably, in prospermatogonia, intergenic H3K4me3 signal intensities were comparable to those at TSSs (Figure 2A; Supplemental Fig. S2E-F), underscoring their potential regulatory importance (Guenther et al. 2007).

**Figure 2.**
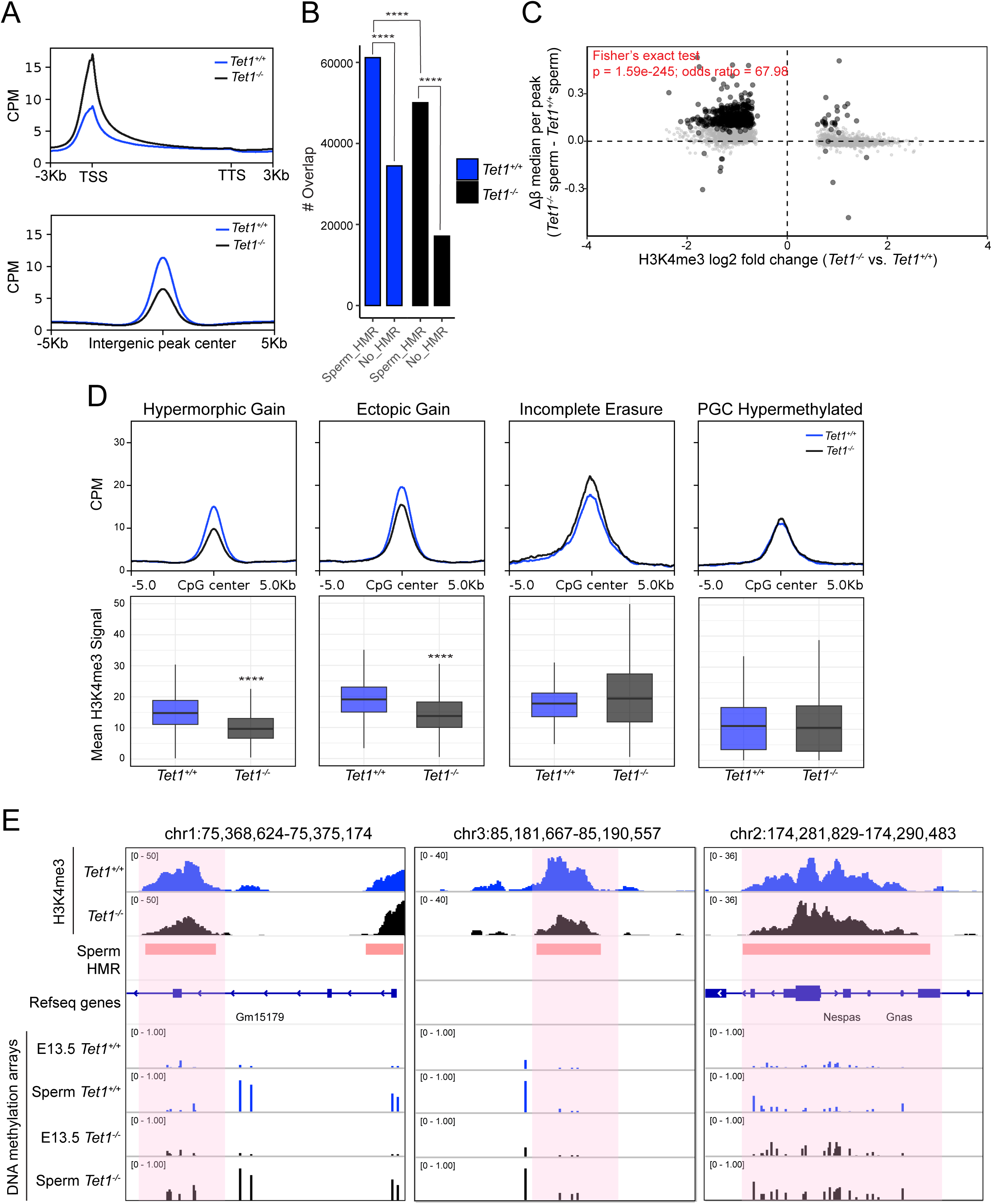
Abnormal de novo methylation acquisition is associated with H3K4me3 erosion in *Tet1^−/−^* male germ cells. A) Metaplot of H3K4me3 signals over genic regions (top) or consensus intergenic peaks (bottom). B) Distribution of H3K4me3 peaks overlapping sperm HMRs compared to non-HMRs (**** p < 0.0001, Fisher’s exact test). C) Scatterplot of *Tet1^−/−^* sperm DMRs and regions with differential H3K4me3 in *Tet1^−/−^*prospermatogonia. H3K4me3 peaks overlapping with DMRs in *Tet1^−/−^* sperm are highlighted in black circles. Fisher’s exact test for hypermethylated DMRs overlapping H3K4me3-depleted vs other regions. D) (Top) Metagene plots of H3K4me3 signals in *Tet1^−/−^* and *Tet1^+/+^*prospermatogonia overlapping subsets of *Tet1^−/−^* sperm hypermethylated DMRs (hypermorphic gain, ectopic gain, incomplete erasure) or *Tet1^−/−^*PGC hypermethylated DMRs. (Bottom) Boxplots of mean H3K4me3 signals over 1 kb windows centered on subsets of hypermethylated DMRs (**** p < 0.0001, Wilcoxon signed-rank test). E) Genome browser tracks for representative sperm hypermethylated DMRs from abnormal de novo methylation at genic region (left) or intergenic region (middle). *Nespas* is a representative ICR that is hypermethylated due to incomplete erasure (right).

To gain further insights into likely genomic targets of TET1 during *de novo* methylation, and their relationship to H3K4me3 changes, we used the 5hmC landscape in wild-type prospermatogonia as a proxy for TET1 localization. As prior work has established that TET antibodies can be an unreliable means of determining TET binding in the absence of added epitope tags (PMID: 25886910; 37456851), localizing 5hmC has been utilized as an alternative proxy for TET binding. Building on these observations, we propose that, even in contexts where its primary function is non-catalytic, TET occupancy can be reliable ascertained by mapping 5hmC (Yan et al. 2022). To that end, we conducted selective chemical labeling of 5hmC followed by affinity enrichment and sequencing (5hmC-Seal) to localize 5hmC in prospermatogonia (Supplemental Fig. S2E-F) (Song et al. 2011). We tested whether regions with decreased or increased H3K4me3 enrichment in *Tet1^−/−^* prospermatogonia were significantly enriched for 5hmC-called peaks compared to random expectation using a permutation-based overlap test (Supplemental Fig. S2G). While regions that gained H3K4me3 did not show significant overlap, H3K4me3 depleted regions were significantly enriched for 5hmC-peaks in prospermatogonia (Supplemental Fig. S2G). As an orthogonal approach, we analyzed publicly available base-resolution maps of 5hmC (APOBEC-seq) for prospermatogonia and defined 5hmC-high regions as 1kb non-overlapping bins where at least 5 CpGs were present with a minimum average of 10% 5hmC across the bins (GSE186357) (Yan et al. 2022). Similar to the correlation between H3K4me3 changes and 5hmC peaks from hmC-Seal, 5hmC-high bins strongly overlapped with H3K4me3 depleted regions in *Tet1^−/−^* prospermatogonia (Supplemental Fig. S2H). These results suggest that erosion of H3K4me3 deposition during *de novo* methylation acquisition in *Tet1^−/−^* prospermatogonia coincides with high likelihood of TET1 localization in WT prospermatogonia. In contrast, the lack of overlap between regions that gained H3K4me3 and 5hmC-high bins implies a less direct relationship to TET1.

As we previously showed, H3K4me3 enrichment in prospermatogonia strongly overlapped with sperm HMRs (Figure 2B, Supplemental Fig. S2D) (Prasasya et al. 2024). With TET1 loss, we observed significantly less overlap (in absolute number of sperm HMRs) between E17.5 H3K4me3 peaks and sperm HMRs (Figure 2B). This initial analysis suggested that fewer sperm HMRs were protected by H3K4me3 in *Tet1^−/−^*prospermatogonia. To follow up, we analyzed the relationship between *Tet1^−/−^*sperm DMRs and changes to H3K4me3 in *Tet1^−/−^*prospermatogonia. With the caveat that the Infinium array provided methylation data for only a representative subset of the genome, we found strong overlap between regions that lost H3K4me3 in *Tet1^−/−^* prospermatogonia and hypermethylated DMRs in *Tet1^−/−^* sperm (Figure 2C). As described above, the majority of *Tet1^−/−^* sperm hypermethylated DMRs originate from abnormal methylation acquisition following successful DNA demethylation in E13.5 PGCs (Figure 1E). As biochemical studies have shown the antagonism between methylated H3K4 and *de novo* DNMTs, we posit that diminished H3K4me3 in *Tet1^−/−^* prospermatogonia increases DNMT3A/3L access, leading to aberrant methylation at these regions. Assessing H3K4me3 signals and *Tet1^−/−^* sperm hypermethylated DMR subsets showed that those originating from abnormal methylation (hypermorphic and ectopic) corresponded to regions with H3K4me3 erosion in prospermatogonia, while those originating from incomplete methylation erasure did not show changes in H3K4me3 signals (Figure 2D). DMRs specific to *Tet1^−/−^* PGCs also showed no H3K4me3 changes at the prospermatogonia stage, suggesting a decoupling between the role of TET1 in DNA demethylation during reprogramming and its role in patterning the chromatin for *de novo* methylation acquisition (Figure 2D). Incidences of abnormal methylation gain that coincide with H3K4me3 reduction in *Tet1^−/−^* prospermatogonia can be found in both genic and intergenic regions (Figure 2E). ICRs such as those from *Nespas/Gnas* locus, where hypermethylation originated from incomplete methylation erasure (Supplemental Fig. S1B), had largely unchanged H3K4me3 in *Tet1^−/−^* prospermatogonia (Figure 2E).

H3K4me3 is highly enriched at TSSs of active genes in eukaryotic cells (Barski et al. 2007; Schneider et al. 2004). The relationship between H3K4me3 and gene transcription is chromatin context dependent, with the strongest evidence demonstrating the need for H3K4me3 for transcriptional elongation but not initiation (Lauberth et al. 2013; Wang et al. 2023). We assessed the relationship between H3K4me3 changes and gene expression in *Tet1^−/−^*prospermatogonia. While total RNA sequencing revealed many differentially expressed transcripts in *Tet1^−/−^* prospermatogonia (DEGs: 2035 upregulated, 4772 downregulated; Supplemental Fig. S3A; Supplemental Table S4A), we did not observe any directional correlations to changes in H3K4me3 (Figure 3A, Supplemental Table S4B). Perhaps consistent with the known promiscuity or leakiness of gene expression in the male germline, where up to 80% of all protein-coding genes can be detected at any given developmental stage, DEGs are associated with biological processes at much later stages of spermatogenesis (e.g. downregulation of motility-associated genes and upregulation of chromosome segregation during meiosis-associated genes; Supplemental Fig. S3B, C) (Xia et al. 2020). Overall, our data showed that H3K4me3 in prospermatogonia does not direct global gene expression.

**Figure 3.**
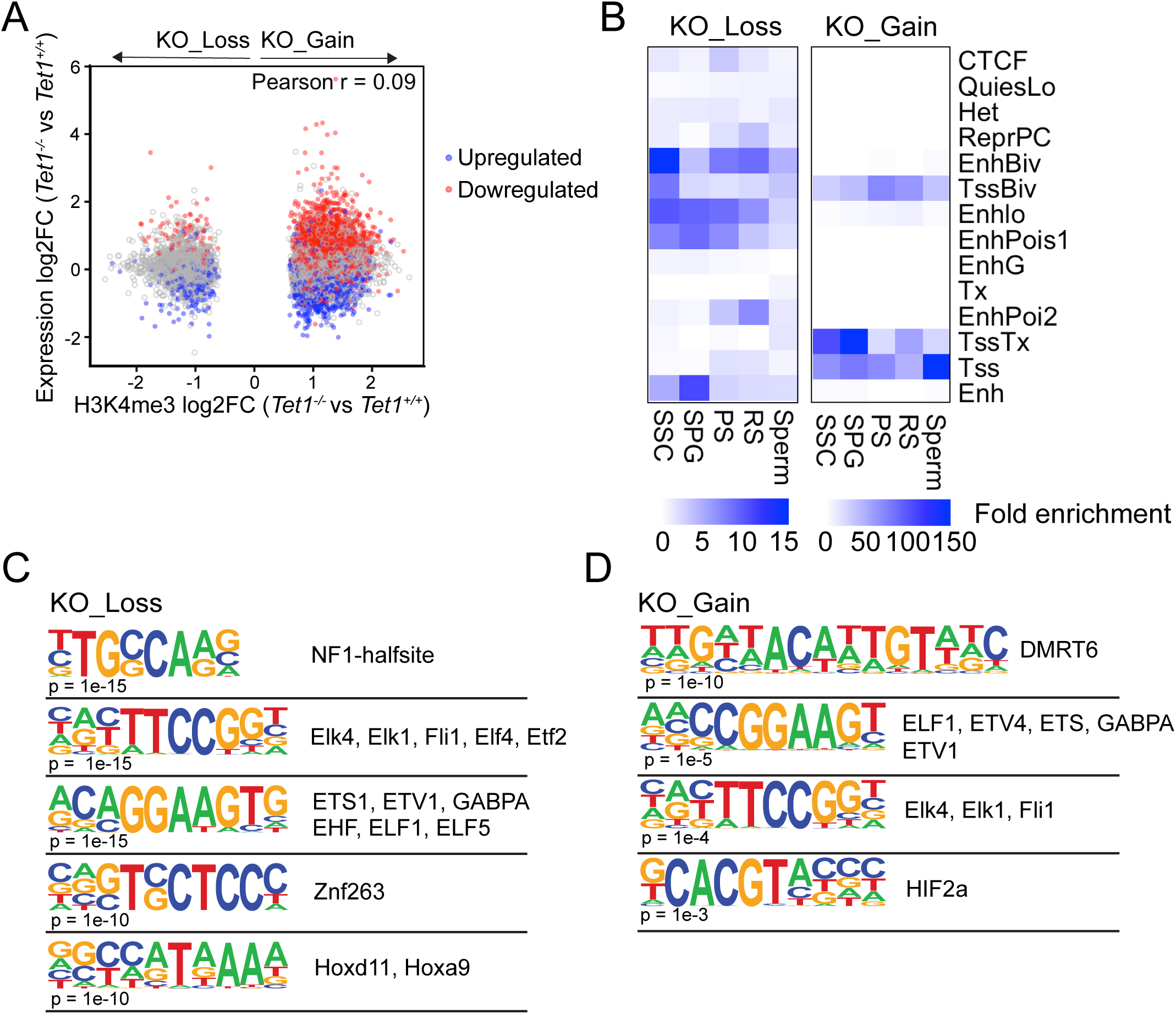
H3K4me3 enrichment in prospermatogonia may regulate later stages of spermatogenesis. A) Scatter plot of significantly altered H3K4me3 peaks in *Tet1^−/−^* prospermatogonia (vs WT) at gene promoters and their associated gene expression changes as measured by total RNA-seq. Differentially expressed transcripts (FDR ≤ 0.05, ≥2-fold change) are highlighted with blue (downregulated) or red (upregulated) dots. B) Enrichment of H3K4me3 peaks lost or gained in *Tet1^−/−^* prospermatogonia across chromatin states during spermatogenesis (SSC: spermatogonial stem cells; SPG: spermatogonia; PS: pachytene spermatocyte; RS: round spermatid), using ChromHMM. Chromatin states assignment is based on the ENCODE consortium projects and are defined in Supplemental Fig. S3D (Ernst and Kellis 2017). Scale bar indicates fold enrichment. C) Motif enrichment for regions with H3K4me3 loss after masking repetitive elements. D) Motif enrichment for regions with H3K4me3 gain.

To explore the regulatory potential of these altered regions across spermatogenesis, we performed chromatin state discovery using ChromHMM. We built segmentation models from publicly available histone PTMs, Pol2, and CTCF ChIP-seq datasets (Supplemental Fig. S3D, Supplemental Table 5) spanning five stages of spermatogenesis: SSCs, differentiated spermatogonia (SPG), pachytene spermatocyte (PS), round spermatid (RS), and sperm. We assigned chromatin states based on Roadmap Epigenomics Consortium conventions (Supplemental Fig. S3D) (Ernst and Kellis 2012). Regions that gained H3K4me3 in *Tet1^−/−^*prospermatogonia matched canonical TSS, while regions with depleted H3K4me3 corresponded to enhancer-like states (Figure 3B). Regions with reduced H3K4me3 in *Tet1^−/−^* prospermatogonia were enriched for long terminal repeats (LTRs) and LINE elements (Supplemental Fig. S3E). Motif enrichment analysis after repeat masking identified recognition sites for methylation sensitive ETS and HOX families of transcription factors (Figure 3C) (Stephens and Poon 2016; Huang et al. 2012). Consistent with potentially serving as promoters that are relevant for male germ line development (Supplemental Fig. S3F), regions that gained H3K4me3 in *Tet1^−/−^* prospermatogonia contain motifs for DMRT6 binding, a germ cell-specific, methylation sensitive transcription factor that coordinates the mitosis to meiosis transition (Figure 3D) (Zhang et al. 2014).

Our findings support the hypothesis that non-catalytic domains of TET1 contribute to H3K4me3 deposition, which may protect intergenic sperm HMRs from *de novo* methylation acquisition. Indeed, emerging evidence shows that non-catalytic domains of TET1 within the C-terminus interact with chromatin modifiers in development and disease (Lian et al. 2016; Joshi et al. 2022). We previously generated a mouse line with the *Tet1^H1654Y,D1656A(HXD)^* mutation, which produces a catalytically inactive but full-length TET1 protein (Ko et al. 2011; Chrysanthou et al. 2022; Caldwell et al. 2021; Prasasya et al. 2024). To test whether non-catalytic TET1 expression is sufficient to establish the chromatin landscape associated with proper male germline *de novo* methylation, we analyzed *Tet1^HxD/HxD^* prospermatogonia.

We conducted CUT&RUN for H3K4me3 in *Tet1^HxD/HxD^*prospermatogonia at E17.5. In support of our hypothesis, the majority of H3K4me3 changes seen in *Tet1^−/−^* prospermatogonia were absent in *Tet1^HxD/HxD^* compared to WT (Figure 4A, Supplemental Fig. S4A, Supplemental Table S2C). Of the regions that lost H3K4me3 in *Tet1^−/−^* prospermatogonia, over 89% (20494 of 22904) were reverted to WT levels in *Tet1^HxD/HxD^* (Figure 4A, Supplemental Fig. S4A, Supplemental Table S3C, D). The genomic distribution of H3K4me3 consensus peaks in *Tet1^HxD/HxD^* prospermatogonia was comparable to *Tet1^−/−^* and WT and consistently overlapped with sperm HMRs (Supplemental Fig. S4B-C). At both TSS-centered and intergenic peaks, H3K4me3 signals in *Tet1^HxD/HxD^* prospermatogonia showed intermediate levels between WT and in *Tet1^−/−^* (i.e. increased around TSSs and decreased at intergenic peaks; Supplemental Fig. S4D-E).

**Figure 4.**
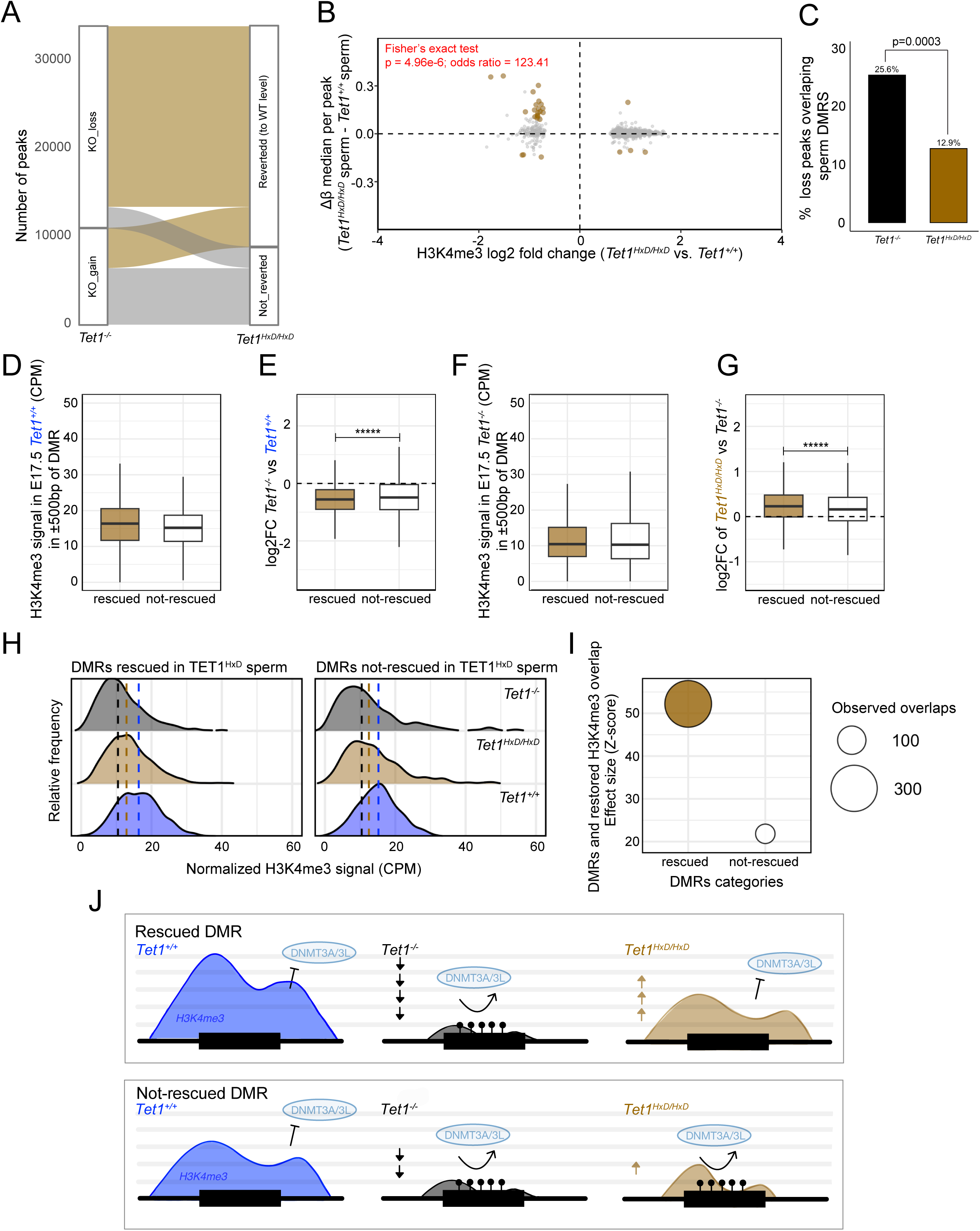
The non-catalytic, full length TET1^HxD^ reverts most H3K4me3 changes in *Tet1^−/−^* prospermatogonia to WT levels. A) Alluvial plot of differentially enriched H3K4me3 peaks in *Tet1^−/−^* vs *Tet1^+/+^* and their status in *Tet1^HxD/HxD^* vs *Tet1^+/+^*. Light brown ribbons indicate peaks altered in *Tet1^−/−^* but not in *Tet1^HxD/HxD^*. B) Scatterplot of *Tet1^HxD/HxD^* sperm DMRs and regions with differential H3K4me3 in *Tet1^HxD/HxD^* prospermatogonia. H3K4me3 peaks overlapping with DMRs in *Tet1^HxD/HxD^* sperm are highlighted in brown circles. Fisher’s exact test for hypermethylated DMRs overlapping H3K4me3-depleted vs other regions. C) Proportion of regions with H3K4me3 loss in Tet1 mutants that overlap hypermethylated sperm DMRs; two-sample proportion test. D) Boxplots of median H3K4me3 signals over 1 kb windows centered on of DMRs that were significantly hypermethylated in *Tet1^−/−^* sperm but were not hypermethylated in *Tet1^HxD/HxD^* sperm (left, rescued DMRs) or stayed hypermethylated in *Tet1^HxD/HxD^* sperm (right, not-rescued DMRs) (Prasasya et al. 2024) (**** p < 0.0001, Wilcoxon signed-rank test). E) Fold change of H3K4me3 signals in *Tet1^−/−^* compared to *Tet1^+/+^* calculated across two subsets of sperm DMRs defined in D (**** p < 0.0001, Wilcoxon signed-rank test). F) Boxplots of median H3K4me3 signals over 1 kb windows centered on hypermethylated DMRs defined in D in *Tet1^−/−^* prospermatogonia (Wilcoxon signed-rank test; not-significant). G) Fold change of H3K4me3 signals in *Tet1^HxD/HxD^* compared to *Tet1^−/−^* calculated across two subsets of sperm DMRs defined in D (**** p < 0.0001, Wilcoxon signed-rank test). H) Ridgeline plots showing the relative frequency distribution of rescued and not-rescued DMRs and the associated H3K4me3 signals found in *Tet1^+/+^*, *Tet1^−/−^*, or *Tet1^HxD/HxD^*prospermatogonia. The median H3K4me3 signal for each genotype across the two subsets of sperm DMRs is indicated with vertical dashed lines of matched colors (blue: *Tet1^+/+^*, black: *Tet1^−/−^*, or brown: *Tet1^HxD/HxD^*). I) The strength of overlap between distinct subsets of DMRs (rescued in *Tet1^HxD/HxD^* sperm or not-rescued in *Tet1^HxD/HxD^* sperm) and regions where H3K4me3 was reverted to WT levels in *Tet1^HxD/HxD^* prospermatogonia following observed depletion in *Tet1^−/−^* prospermatogonia as determined by permutation test (Z-score from 1000 permutations of randomly selected regions consistent with Illumina Bead Chip array coverage). J) Schematic diagram depicting differential responsiveness in the relative levels of H3K4me3 depletion in response to the loss of TET1 (*Tet1^⁻/⁻^*, black, compared to *Tet1^+/+^*, blue). This depletion correlates with aberrant methylation acquisition, resulting in hypermethylated DMRs in *Tet1^⁻/⁻^* sperm. Catalytically inactive TET1^HxD^ enhances H3K4me3 deposition in prospermatogonia toward WT levels. However, not all restoration of H3K4me3 is associated with rescue of abnormal methylation. It is plausible that additional chromatin features, beyond H3K4me3, may contribute to differential susceptibility to aberrant *de novo* methylation at the two DMR subsets.

The limited number of regions where H3K4me3 was depleted in *Tet1^HxD/HxD^*prospermatogonia (compared to WT) remained associated with eventual hypermethylated DMRs in *Tet1^HxD/HxD^* sperm, although the overlap was significantly reduced compared to *Tet1^−/−^*germ cells (Figure 4B-C). Among 5631 hypermethylated DMRs identified in *Tet1^−/−^*sperm, 3710 (∼2/3) were not significantly hypermethylated in *Tet1^HxD/HxD^* sperm (defined as rescued DMRs)(Prasasya et al. 2024). These methylation defects cannot be fully attributed to loss of TET1 catalytic activity. Of note, regions that remained hypermethylated in *Tet1^HxD/HxD^* sperm are those that require TET1-iterative oxidation capability such as ICRs and promoter of germline responsive genes (Prasasya et al. 2024).

We examined how H3K4me3 enrichment in prospermatogonia relates to the rescue of sperm hypermethylation defects in *Tet1^HxD/HxD^* compared to *Tet1^−/−^* sperm. First, we assessed the baseline levels of H3K4me3 in WT prospermatogonia for rescued and not-rescued DMRs and found rescued DMRs display modestly higher baseline levels (Figure 4D). Following the loss of TET1, rescued DMRs underwent a greater depletion of H3K4me3 compared to not-rescued DMRs (Figure 4E), resulting in the convergence of H3K4me3 levels between the two subsets in *Tet1^−/−^* prospermatogonia (Figure 4F). To quantify the extent of H3K4me3 restoration toward WT levels, we calculated the fold-change of H3K4me3 signal in *Tet1^HxD/HxD^* relative to *Tet1^−/−^*prospermatogonia. The degree of restoration was significantly greater across rescued DMRs than across those that remained hypermethylated (not-rescued; Figure 4G). Ridgeline plots in Figure 4H demonstrate the difference in dynamic range of H3K4me3 regulation in WT, *Tet1^−/−^*, and *Tet1^HxD/HxD^* between the two DMR subsets. Overall, rescued DMRs exhibit greater susceptibility to H3K4me3 depletion following TET1 loss, but also greater responsiveness in H3K4me3 restoration in the presence of non-catalytic TET1^HxD^.

To test this relationship more formally, we performed permutation-based enrichment analysis to assess whether rescued DMRs preferentially overlapped regions showing reverted H3K4me3 to WT levels in *Tet1^HxD/HxD^* prospermatogonia (Gel et al. 2015). Both rescued and not-rescued DMRs showed significant enrichment within H3K4me3-reverted peaks in *Tet1^HxD/HxD^*prospermatogonia (Supplemental Fig. S4F-G), but the effect size (Z-score) was markedly higher for rescued DMRs (Figure 4I). Together, these results indicate that non-catalytic full length TET1^HxD^ is sufficient to enhance H3K4me3 deposition overall in prospermatogonia compared to the absence of TET1. Taken together, our findings suggest that the susceptibility of a region to aberrant DNMT3A/3L reflects the extent, and likely timing, of H3K4me3 erosion in *Tet1^−/−^*prospermatogonia during the *de novo* methylation window (illustrated in Figure 4J).

In the male germline, hypermethylation is required, in part, to silence retroelements that are toxic for meiosis (Bourc’his and Bestor 2004; Barau et al. 2016). Despite this, certain regions of the mammalian sperm genome resist DNA methylation, including CpG-dense promoters and younger, species-specific transposon subfamilies overlapping regulatory elements (Shirane and Lorincz 2023; Molaro et al. 2011; Erkek et al. 2013). Additionally, low methylation at CpG-dense regions correlates with nucleosome retention during histone-to-protamine exchange, a critical step male gamete maturation (Erkek et al. 2013). Retained nucleosomes in sperm play a crucial role in patterning chromatin of the early embryo, underscoring the importance of the sperm epigenome beyond spermatogenesis (Fanourgakis et al. 2025). Thus, the integrity of sperm HMRs is essential for both spermatogenesis and potentially embryo viability.

We previously demonstrated that most methylation defects from compromised TET1 activity occur within sperm HMRs (Prasasya et al. 2024). Iterative oxidation is required at ICRs and meiosis-associated promoters, but these account for only a small fraction of *Tet1^−/−^* sperm defects. In addition to its role in active demethylation, TET1 has been proposed to protect against DNA re-methylation during epigenetic reprogramming (Hill et al. 2018). However, DNMT3A/3L activity is limited at E14.5, making this role less likely in PGCs. Our findings instead suggest a role for TET1 in preventing abnormal methylation acquisition in prospermatogonia. Aberrant DNA methylation in *Tet1^−/−^* sperm correlates with erosion of H3K4me3 at the prospermatogonia stage, a chromatin feature that negatively correlates with hypermethylated regions in the male germline and a mark that has been biochemically demonstrated to be the most antagonistic to DNA methyltransferase access (Figure 4J). Consistently, the non-catalytic function of TET1, as tested in *Tet1^HxD/HxD^* germline, is sufficient to enhance H3K4me3 deposition and partially rescue DNA methylation (Prasasya et al. 2024). Analysis of H3K4me3 enrichment in *Tet1^HxD/HxD^* prospermatogonia suggests that there are differential dependencies – likely related to both the magnitude and timing of H3K4me3 changes – that influence TET1 non-catalytic functions and antagonism of DNMT3A/3L during *de novo* methylation, as not all regions with H3K4me3 restoration corresponds to rescue of DNA hypermethylation (Figure 4J). It is notable that our model only allows for correlative interpretations for the importance of H3K4me3 in shaping the sperm HMRs landscape. Additional studies to conditionally target H3K4 methyltransferases during *de novo* methylation period are needed to determine the governing rule by which DNMT3A/3L is excluded from sperm HMRs.

TET proteins interact with chromatin modifiers in many contexts. In embryonic stem cells (ESCs), the non-catalytic function of TET1 shapes bivalency (H3K4me3, H3K27me3) at regulators of mesoderm and trophectoderm lineages by recruiting Polycomb repressive complex 2 (PRC2) and Sin3a (Chrysanthou et al. 2022). TET1 is also a stable binding partner of *O-*GlcNAc transferase (OGT) and is required for OGT-Sin3a-Hcf1 binding to chromatin (Vella et al. 2013; Williams et al. 2011). Hcf1 is a key component of the SET1/COMPASS histone methyltransferase complex, a writer for H3K4me2/3. In hematopoiesis, the TET2/3-OGT complex promotes Hcf1 O-GlcNAcylation, leading to SET1/COMPASS assembly, H3K4me3 deposition, and gene activation (Capotosti et al. 2011; Deplus et al. 2013). A similar mechanism is plausible in prospermatogonia, as single cell RNA-seq confirms *Ogt* and *Hcfc1* expression in fetal and neonatal germ cells, coinciding with *de novo* methylation (Zhao et al. 2021). TET1 recruitment to regions with TET1-dependent H3K4me3 deposition in prospermatogonia is further supported by the strong overlap between 5hmC localization and H3K4me3 depleted regions in *Tet1^−/−^* prospermatogonia as shown by the finding of 5hmC-Seal analysis (Supplemental Fig. S2G).

It is also possible that in addition to promoting H3K4me3 deposition, TET1 may also partially restrain DNA methylation by physically protecting its targets from DNMT3A/3L in prospermatogonia. This is not likely to be the main mechanism of sperm HMRs protection however, as TET1 and DNMT3A antagonism was previously found to be highly context dependent within proximal promoters (Gu et al. 2018). Furthermore, in human ESCs, TET1 and DNMT3A have been shown to co-bind, especially within DNMT3A preference binding regions (Chao et al. 2022). Notably, TET1-dependent sperm HMRs are intergenic and acquire enhancer-like chromatin states throughout spermatogenesis (Figure 2A, 3B). Top enriched motifs in these regions correspond to ETS transcription factors. Significantly, the ETS family of transcription factors are known to interact with and recruit histone acetyltransferases such as CBP and p300, in further support of the potential of intergenic sperm HMRs as regulatory elements (Yang et al. 1998; Goodman and Smolik 2000). We propose that protecting intergenic sperm HMRs from methylation helps establish the enhancer landscape that regulates stage-specific gene expression, particularly during the mitosis-to-meiosis transition (Maezawa et al. 2020). Indeed, *Tet1^−/−^* male mice reproductive defects are attributed to premature aging due to reduced SSCs and progressively worse meiotic defects (Huang et al. 2020). While the mechanism orchestrating chromatin patterning before *de novo* methylation remains unclear, our findings reveal the non-catalytic function of TET1 as a molecular component for establishing sperm HMRs.

## Methods

### Mouse models and germ cell collection

All experiments were approved by the Institutional Animal Care and Use Committee of the University of Pennsylvania (protocol number: 804211). Mice were housed in polysulfone cages within a pathogen-free facility under a 12 h light/dark cycle, with ad libitum access to water and standard chow (Laboratory Autoclavable Rodent Diet 5010, LabDiet, St. Louis, MO, USA). *Tet1* knockout mice (Dawlaty et al. 2013) (017358; B6;129S4-Tet1^tm1.1Jae^/J) and *Oct4-GFP* mice (Lengner et al. 2007) (008214; B6;129S4-Pou5f1^tm2Jae^/J) were obtained from The Jackson Laboratory and backcrossed for at least 10 generations to a C57BL/6J background (000664; B6; The Jackson Laboratory). Catalytically inactive *Tet1^HxD^* mouse line was generated as described in (Prasasya et al. 2024). The *Oct4-GFP* allele was maintained as homozygous in *Tet1* heterozygote breeders (*Tet1^+/−^*; *Oct4^GFP/GFP^* or *Tet1^HxD/+^; Oct4^GFP/GFP^)*. Primers to determine sex, *Tet1^tm1.1Jae^*, *Tet1^HxD^*, and *Oct4-GFP* alleles are listed in Supplemental Table S6.

Gonadal tissues from E13.5 and E17.5 embryos (post copulatory plug detection) were dissected and digested into a single-cell suspension using 0.25% trypsin/EDTA for 10 minutes at 37 °C. *GFP^+^*germ cells were isolated by flow cytometry. For bisulfite mutagenesis and RNA isolation, germ cells were sorted into empty tubes and immediately snap-frozen for storage at −80 °C until processing. For CUT&RUN, sorted cells were collected into chilled PGC medium (DMEM supplemented with 15% FBS, 100 U/mL leukemia inhibitory factor, 5 µM forskolin, 1 ng/mL stem cell factor). Following sorting, dimethyl sulfoxide (DMSO) was added to a final concentration of 10%, and the cells were slow frozen using Mr. Frosty until library preparation.

### Genome-wide DNA methylation profiling using Infinium Mouse Methylation BeadChip

Snap frozen germ cell pellet was directly lysed and bisulfite mutagenized using EZ DNA Methylation Direct Kit (Zymo Research) per manufacturer’s instructions. Bisulfite mutagenized (BSM) DNA was subjected to low-input whole-genome amplification workflow prior to hybridization to Illumina BeadChip array as detailed in (Lee et al. 2024) (Workflow J). Briefly, BSM DNA underwent 2 rounds of Klenow amplifications (NEB) using random hexamer primers (NEB). For additional details on library preparation and data analysis, see the Supplemental Material. Amplified DNA was cleaned using solid-phase reversible immobilization (SPRI) beads (2x; Beckman Coulter) and concentration was measured with Qubit (Invitrogen).10 µL of BSM DNA library was submitted for hybridization onto the Infinium Mouse Methylation-12v1-0 BeadChip (Illumina) at Children’s Hospital of Philadelphia Center for Applied Genomics.

### CUT&RUN library preparation

CUT&RUN experiments were performed on samples of 4,000 nuclei isolated per epitope from cryopreserved germ cells as described above using CUTANA ChIC/CUT&RUN kit (EpiCypher). Three biological replicates per genotype were prepared for H3K4me3 (1:100, Cell Signaling Technology antibody 2729) and IgG (1:50, CUTANA Rabbit IgG negative control antibody). Cells were rapidly thawed and counted, followed by nuclei extraction using NE buffer (20 mM HEPES-KOH pH 7.9, 10mM KCl, 0.5 mM fresh spermidine, 0.1% Triton X-100, 20% glycerol plus protease inhibitors). Nuclei were bound to concanavalin A-coated magnetic beads and CUT&RUN was conducted per CUTANA product manual with modifications. CUT&RUN libraries were prepared with NEBNext Ultra II DNA Library Prep Kit for Illumina (NEB). For additional details on library preparation and data analysis, see the Supplemental Material.

### Total RNAseq library preparation

Total RNA was extracted from 5000 snap frozen E17.5 germ cells isolated from a single embryo using TRIzol reagent (Invitrogen). RNA integrity number and concentration were assessed using the High Sensitivity RNA ScreenTape for TapeStation system (Agilent Technologies). Ribosomal RNA-depleted total RNA libraries were generated using the SMART-Seq Total RNA Pico Input Kit with UMIs with ZapR Mammalian (Takara Bio) from 5 ng input according to the manufacturer’s protocol. Libraries were indexed with NEBNext Multiplex Oligos for Illumina Unique Dual Index Primer Pairs (NEB) and were amplified using SeqAmp DNA Polymerase (Takara Bio). For additional details on library preparation and data analysis, see the Supplemental Material.

### 5hmC-Seal library preparation

Genomic DNA was isolated from 30,000 snap frozen E17.5 germ cells pooled from multiple embryos using Phenol:Chloroform:Isoamyl Alcohol (25:24:1; Sigma-Aldrich) and ethanol precipitated, followed by resuspension in 10 mM Tris-HCl pH 8.0. 50 ng of genomic DNA was sheared using NEBNext UltraShear (NEB) and was used as input for enrichment of 5-hydroxymethylcytosine using 5hmC-Seal following manufacturer protocol (5hmC Profiling Kit; Active Motif). Sequencing libraries were prepared from both input and enriched fractions using NEBNext UltraII DNA Library Prep Kit for Illumina (NEB).

### Data availability

The accession number for raw and processed Illumina Mouse Infinium BeadChip, CUT&RUN, RNA-seq, and 5hmC-Seal data generated in this paper is submitted to GEO with the accession number GSE318256. Any additional information required to reanalyze the data reported in this paper is available from corresponding author upon request.

## Extended Methods

### Genome-wide amplification of bisulfite mutagenized DNA

10 µL of bisulfite mutagenized (BSM) DNA eluted from EZ DNA methylation-Direct Kit (Zymo Research) was combined with 0.8 µL 10 mM dNTPs (BioRad, F560/170-8874), 4 µL 20 µM Random Primer 6 (NEB, S1230S), and nuclease-free water to a final volume of 16 µL. Samples were denatured at 98 °C for 1 min, rapidly cooled on ice for 2 min, then incubated at 37 °C for 30 min with 2 µL (10 U) of Klenow Fragment (3′→5′ exo–; NEB, M0212S) and 2 µL of the supplied buffer. A second round of Klenow amplification was done by denaturing samples for 98°C for 1 min, followed by rapid cooling on ice. 2 µL of Klenow Fragment, 0.5 µL of supplied buffer, 0.8 µL of dNTPs, and 1.7 µL of water were added, followed by 37°C incubation for 30 minutes. Reactions were heat-inactivated at 75 °C for 10 min and purified with SPRI beads (Beckman Coulter) at a 2× bead-to-sample volume ratio. Libraries were eluted in 11 µL nuclease-free water and DNA concentration was measured using Qubit High Sensitivity DNA Reagent (Invitrogen).

Libraries were denatured with 0.1 N NaOH immediately prior to hybridization onto the Infinium Mouse Methylation-12v1-0 BeadChip (Illumina). TECAN automation was used for sample loading onto the iScan System. Biological replicates for each genotype were as follows: E13.5, *Tet1^+/+^* (n = 8), *Tet1^−/−^* (n = 7).

### CUT&RUN and Sequencing library preparation

CUT&RUN experiments were performed on samples of 4,000 nuclei isolated per epitope from cryopreserved germ cells as described above using CUTANA ChIC/CUT&RUN kit (EpiCypher). Cells were rapidly thawed and counted, followed by nuclei extraction using NE buffer (20 mM HEPES-KOH pH 7.9, 10mM KCl, 0.5 mM fresh spermidine, 0.1% Triton X-100, 20% glycerol plus protease inhibitors). Nuclei were bound to concanavalin A-coated magnetic beads pre-washed in cold bead activation buffer (EpiCypher) for 10 minutes at room temperature (RT). Nuclei were resuspended with H3K4me3 (1:100, Cell Signaling Technology antibody 2729) or IgG (1:50, CUTANA Rabbit IgG negative control antibody) in 50 µL of antibody binding buffer (CUTANA pre-wash buffer – 20 mM HEPES pH 7.5, 150 mM NaCl-, with 0.5 mM spermidine, 1x Roche complete protease inhibitor, 0.05% digitonin and 2mM EDTA) overnight at 4°C. Samples were washed twice in cell permeabilization buffer (CUTANA pre-wash buffer with 0.5mM spermidine, 1x Roche complete protease inhibitor, and 0.05% digitonin) followed by incubation with pAG-MNase fusion protein for 1 hour at 4°C. Cells were washed twice in cell permeabilization buffer, and 100mM CaCl_2_ was added to activate cleavage for 1 hour at 4°C. 2x CUTANA STOP buffer (170 mM NaCl, 20 mM EGTA, 0.05% digitonin, 50 µg/mL RNase A, 25µg/mL Glycogen) was added to quench the reaction. To release antibody-bound fragments, samples were incubated at 37 °C for 10 min, and the supernatant was isolated from the magnet-bound beads. 3 µL of 10% SDS and 2.5 µL of Proteinase K (20 mg/mL) were added to the samples (300 µL in 0.1x TE) and incubated at 70 °C for 10 min to digest residual proteins. DNA was extracted twice with phenol:chloroform with additional glycogen during DNA precipitation. Pellets were resuspended in 50 µL of Tris-EDTA buffer (1 mM Tris-HCl, pH 8; 0.1 mM EDTA).

CUT&RUN and 5hmC-Seal libraries were prepared using the NEBNext Ultra II DNA Library Prep Kit for Illumina with modifications. 1:25 fold dilution of Illumina adapter was used during adaptor ligation and libraries were amplified for 12–14 cycles using NEBNext Multiplex Oligos for Illumina Unique Dual Index Primer Pairs. Amplified libraries were cleaned using solid-phase reversible immobilization (SPRI) beads (Beckman Coulter), quantified using the Kapa library quantification kit for Illumina, and assessed for size distribution using the D5000 ScreenTape Assay for TapeStation System (Agilent Technologies). Libraries were sequenced on Illumina NextSeq1000 instrument (60bpx60bp paired end).

### RNA isolation

Snap-frozen cell pellets were directly transferred to 500 µL TRIzol Reagent (Invitrogen). Samples were incubated at RT for 10 min, followed by addition of 100 µL chloroform and incubation at RT for 3 min. Phase separation was performed by transferring mixture to MaXtract High Density tube (Qiagen) and centrifuging at maximum speed for 10 min at 4 °C. The aqueous phase was collected, mixed with 500 µL isopropanol and 2 µL of 2 mg/mL glycogen, and incubated at RT for 10 min to precipitate RNA. Samples were centrifuged at maximum speed for 10 min at 4 °C. The supernatant was discarded, and the RNA pellet was washed with 80% ethanol, briefly spun to eliminate residual ethanol, and air-dried for 5 min. RNA was resuspended in 17.5 µL TE buffer. To eliminate carry-over genomic DNA 2 µL Qiagen Buffer RDD and 0.5 µL Qiagen DNaseI were added and incubated at RT for 10 min. RNA was cleaned using 1.8× RNAClean XP Beads (Beckman Coulter) and RNA was eluted in 8 µL TE buffer.

### Methylation array data analysis

Raw IDAT files were processed using SeSAMe R Package and the MM285 array manifest file to obtain methylation β-values with prep=”I” (Zhou et al. 2018; Lee et al. 2024). Between-run batch correction was done using the R Package limma removeBatchEffect(). To determine differentially methylated probes, we included only CG probes with no NA values in all biological replicates. Batch-corrected β-values were converted into M-values (log2 ratio of methylated/unmethylated intensities). We then fit a linear model across conditions using the R package limma lmFit() function (Ritchie et al. 2015). We applied eBayes() to moderate the per-locus variances and improved statistical power by borrowing information across loci. The contrast of interest (*Tet1^−/−^* vs *Tet1^+/+^* PGCs) was extracted and significance was determined using FDR (Benjamini-Hochberg) ≤ 0.05 and a minimum effect size of abs(delta. β) > 0.10. We used SeSAMe knowYourCG tool to test enrichments for probe design groups. A differentially methylated region (DMR) corresponds to each differentially methylated probe. Average β values (from biological replicates) of DMRs found in *Tet1^−/−^* PGCs and *Tet1^−/−^* sperm were used to create distribution plot to identify origin of hypermethylation.

### Identification of sperm hypomethylated regions

Sperm hypomethylated regions (HMRs) were determined using hmr function of DNMTools, which uses hidden Markov model approach to identify methylation canyon in a supplied whole genome bisulfite sequencing methylation call data set (GSE56697: Wang et al. 2014).

### ChromHMM model of methylation

The ChromHMM genome segmentation model was built using CTCF, Pol2, H3K4me1/2/3, H3K9me3, H3K27me3, H3K36me3, H3K9ac and H3K27ac datasets from spermatogonial stem cell (SSC), differentiated spermatogonia (SPG), pachytene spermatocytes (PS), round spermatids (RS), and sperm (GSE42629: Erkek et al. 2013; GSE158360: Chen et al. 2021; GSE79227: Jung et al. 2017; GSE89502: Maezawa et al. 2018; GSE207325: Pepin et al. 2024; GSE229246: Fanourgakis et al. 2025; GSE55471: Siklenka et al. 2015; GSE79227: Jung et al. 2017; GSE97778: Wang et al. 2018; GSE132446: Chen et al. 2020; GSE130652: Maezawa et al. 2020; GSE244675: Kitamura et al. 2023; GSE132054: Vara et al. 2019; GSE232398: Kang et al. 2025; GSE108717: Zuo et al. 2018; GSE153063: Yin et al. 2021; GSE49621: Hammoud et al. 2014; GSE137744: Liu et al. 2019; GSE132054: Vara et al. 2019; GSE44346: Margolin et al. 2014; GSE131656: Cheng et al. 2020). BAM files were binarized using the BinarizeBam function from ChromHMM with a Poisson threshold of 0.00001 and other default parameters. State models with 4–16 states were built using the LearnModel function with default parameters. 14 states were selected for the final model. Chromatin state assignment is based on ENCODE consortium convention (Ernst and Kellis 2017).

### RNAseq data analysis

Unique molecular index sequences were located in adaptor positions from raw reads using UMI-tools (Smith et al. 2017). UMI sequences were added to the read names and reads were quality-trimmed using Trim Galore (--phred 33 –clip_R2 6). Reads were aligned to mouse mm10 reference genome using STAR with default parameters and a maximum fragment size of 2000bp (Dobin et al. 2013). Reads with the same mapping coordinates and identical UMIs were collapsed to a single read.

Properly paired primary alignments were retained for downstream analysis using Samtools. Count matrices were generated using featureCounts against GENCODE annotation release M25 (Liao et al. 2014). To remove lowly expressed genes prior to differential expression analysis, we applied library-size aware filtering using the R package edgeR filterByExpr() function. This approach retains genes with at least ∼10 raw counts, scaled to each library’s sequencing depth, in a sufficient number of replicates within at least one experimental group. For our libraries, this corresponded to CPM cutoffs ranging from ∼0.5 to 0.8 across samples. Filtered count matrix were fed into the R package DESeq2 to perform normalization and differential expression analysis with FDR cut off ≤ 0.05 and minimum fold change of 2.

### CUT&RUN and 5hmC-Seal data analysis

Raw reads were quality trimmed using Trim Galore (https://www.bioinformatics.babraham.ac.uk/projects/trim_galore/) using default parameters. Trimmed reads were aligned to mouse mm10 reference genome using Bowtie2 with parameters: –end-to-end –very-sensitive (Langmead and Salzberg 2012). Following alignment, low quality reads (QMAP ≤ 10) and non-primary alignments were removed using SAMtools (view -q 10 -F 256) (Danecek et al. 2021). Duplicated reads were removed using SAMtool rmdup with default parameters and mitochondrial reads were removed using grep function. Bigwig files normalized to count per million (CPM) were prepared using deepTools bamCoverage function with parameters: -bs 1 -normalizeUsing CPM (Ramírez et al. 2016). Peak calling was performed using Model-based Analysis of ChIP-seq (macs2; -qvalue 0.01, narrowPeak setting for CUT&RUN; broadPeak setting for 5hmC-Seal) against sample-matched IgG library as the background (Zhang et al. 2008). Peak sets and aligned reads were fed into the R Bioconductor package DiffBind where consensus peaks were determined based on the presence of peaks in at least two out of three biological replicates. DiffBind was used to perform normalization and differential enrichment analysis with FDR cut-off of ≤ 0.05 and minimum fold change of 1.5. Consensus or differentially enriched peaks were annotated for transcriptional start sites (TSS: −1kb to +100bp), exons, introns, transcriptional termination site (TTS: −100bp to +kb), or intergenic regions using HOMER annotatePeaks.pl function based on GENCODE annotation release M25 (Heinz et al. 2010).

For visualization, average bigwig files over the three replicates were created using bigwigCompare from deepTools. Metagene plots and heatmaps were generated using the computeMatrix, plotProfile, or plotHeatmap from deepTools with parameters reference point b-5000 a-5000 for peak centers or -beforeRegionStartLength 3000 -afterRegionStartLength 3000 for genic regions. Overlap between H3K4me3 signals and *Tet1^−/−^* sperm differentially methylated regions (DMRs) of differing origins were identified using the R bioconductor package GenomicRanges findOverlaps function. Pairwise Wilcoxon rank sum test was used to compare average H3K4me3 signals across the overlap hits. Overlap between differentially enriched H3K4me3 peaks in *Tet1^−/−^* prospermatogonia and all *Tet1^−/−^* sperm DMRs were identified using findOverlaps function. Fisher’s exact test was used to test whether *Tet1^−/−^* sperm DMRs were significantly found within regions that lost H3K4me3. H3K4me3 fold changes of *Tet1^−/−^* vs WT or *Tet1^HxD/HxD^* vs WT were overlapped with TET1^HxD^-rescued hypermethylated DMRs as defined in Prasasya et al. 2024 (FDR ≥ 0.05 in *Tet1^HxD/HxD^*vs WT; FDR ≤ 0.05, delta.beta > 0.10 in *Tet1^−/−^*vs WT). Pairwise Wilcoxon rank sum test with continuity correction was used to compare closeness to WT, which is calculated over fold change of H3K4me3 signals in *Tet1^HxD/HxD^* vs WT compared to *Tet1^−/−^* vs WT across rescued- or not-rescued DMRs.

To test the significance of overlaps between rescued or not-rescued subsets of DMRs with H3K4me3 regions that were restored in *Tet1^HxD/HxD^*compared to *Tet1^−/−^,* we conducted 1000 permutations using R package regioneR. The test was implemented with permTest() using 1000 randomizations where at each iteration, the query regions were randomly shuffled across CpGs covered within the Illumina Bead Chip array while preserving their number and size distribution. For each randomized set, the overlap with the reference feature set was recalculated to generate a null distribution of expected overlaps. The empirical p-value was estimated as the proportion of permutations with equal or more extreme overlaps than observed.

R package ggplot2 was used to create box-and-whisker plots, distribution plots, scatter plots, dot plots, and alluvial plot. R package pheatmap was used to create heatmap. R package BioVenn was used to create venn diagrams.

### Data availability

The accession number for raw and processed Illumina Mouse Infinium BeadChip, CUT&RUN, RNA-seq, and 5hmC-Seal data generated in this paper is submitted to GEO with the accession number GSE318256.

---

**Supplemental Figure 1.**
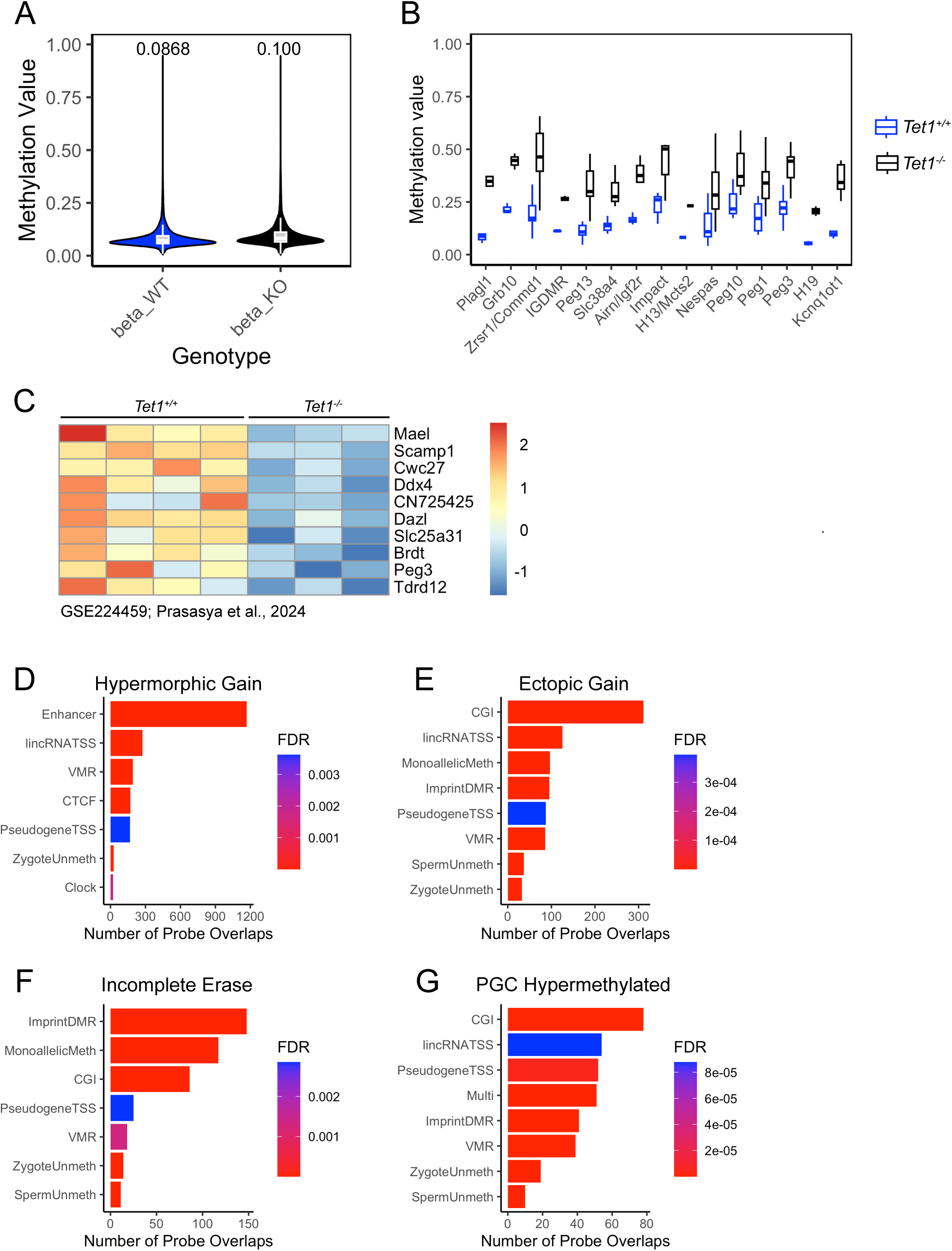
Additional analysis of methylation arrays for Tet1−/− vs Tet1+/+ PGCs. A) Violin plots of average methylation values at each CpG probe with overall mean methylation indicated (n=8 per group). Differentially methylated probes were identified using limma (FDR ≤ 0.05, >10% difference). B) ICRs are the most enriched design-group category among hypermethylated DMRs in *Tet1^−/−^*PGCs (15 of 21 array-represented ICRs are significantly hypermethylated). Boxplots show average methylation of probes associated with hypermethylated ICRs. C–F) Top enriched Infinium Mouse Methylation BeadChip design groups for different categories of *Tet1^−/−^* sperm and PGC DMRs.

**Supplemental Figure 2.**
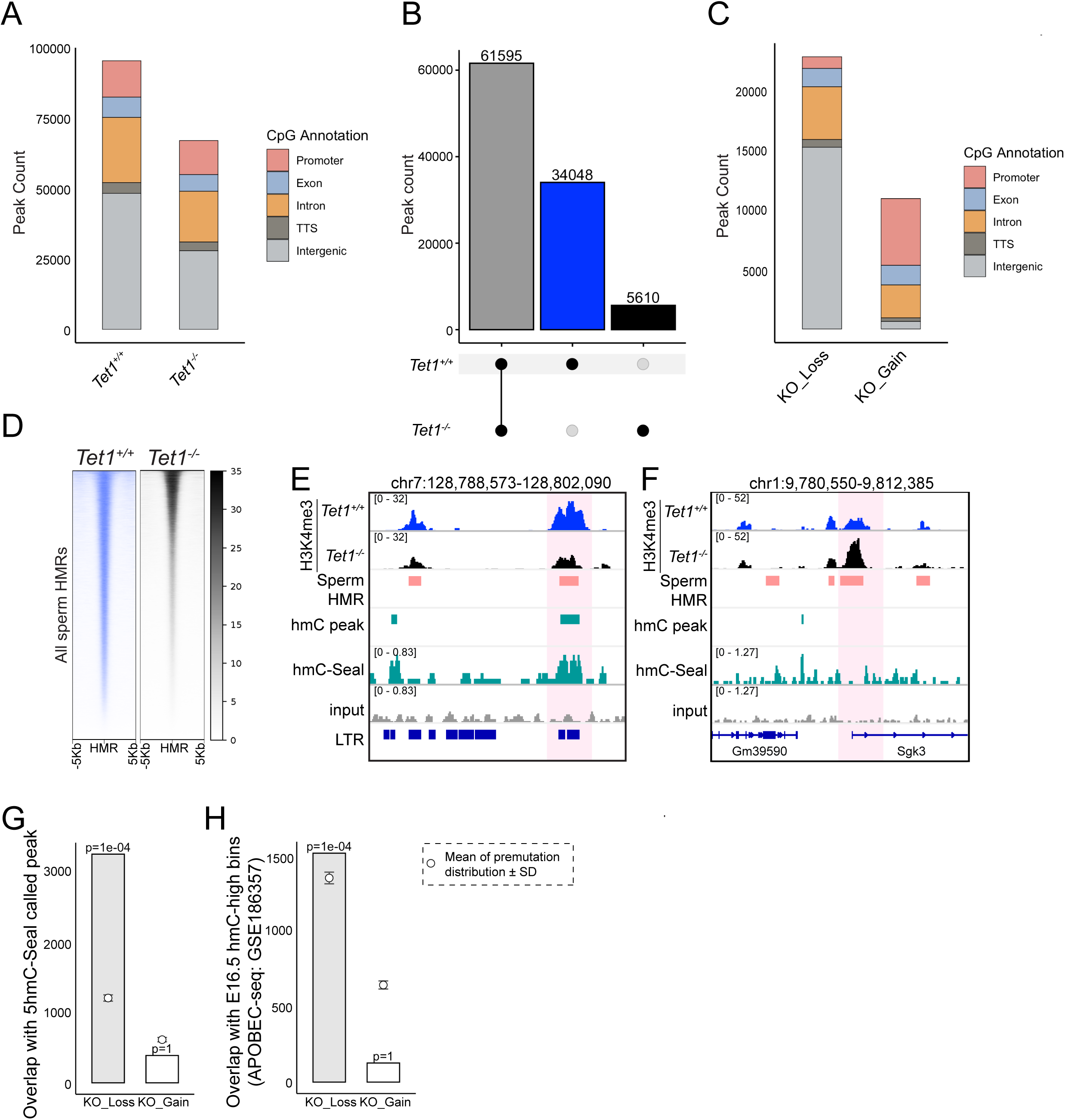
Additional analysis of differentially enriched H3K4me3 signals in *Tet1^−/−^* vs *Tet1^+/+^* prospermatogonia. A) Genomic compartmentalization of H3K4me3 consensus peaks (present in ≥2 of 3 replicates) in *Tet1^+/+^* and *Tet1^−/−^* prospermatogonia. B)) Distribution of H3K4me3 consensus peaks unique to or shared between *Tet1^+/+^* (n=3) and *Tet1^−/−^* (n=3) E17.5 prospermatogonia detected by CUT&RUN. C) Heatmaps of H3K4me3 signals in E17.5 *Tet1^−/−^* and *Tet1^+/+^* prospermatogonia overlapping sperm hypomethylated regions (HMRs) defined by sperm WGBS (GSE56697; Wang et al. 2014). Plots represent averages of 3 biological replicates. D) Genomic distribution of differentially enriched H3K4me3 peaks between *Tet1^−/−^* and *Tet1^+/+^* prospermatogonia (DiffBind, FDR ≤ 0.05, ≥1.5-fold change). E-F) Genome browser tracks for representative regions showing H3K4me3 loss (E, intergenic) or gain (F, promoter) in *Tet1^−/−^*prospermatogonia.

**Supplemental Figure 3.**
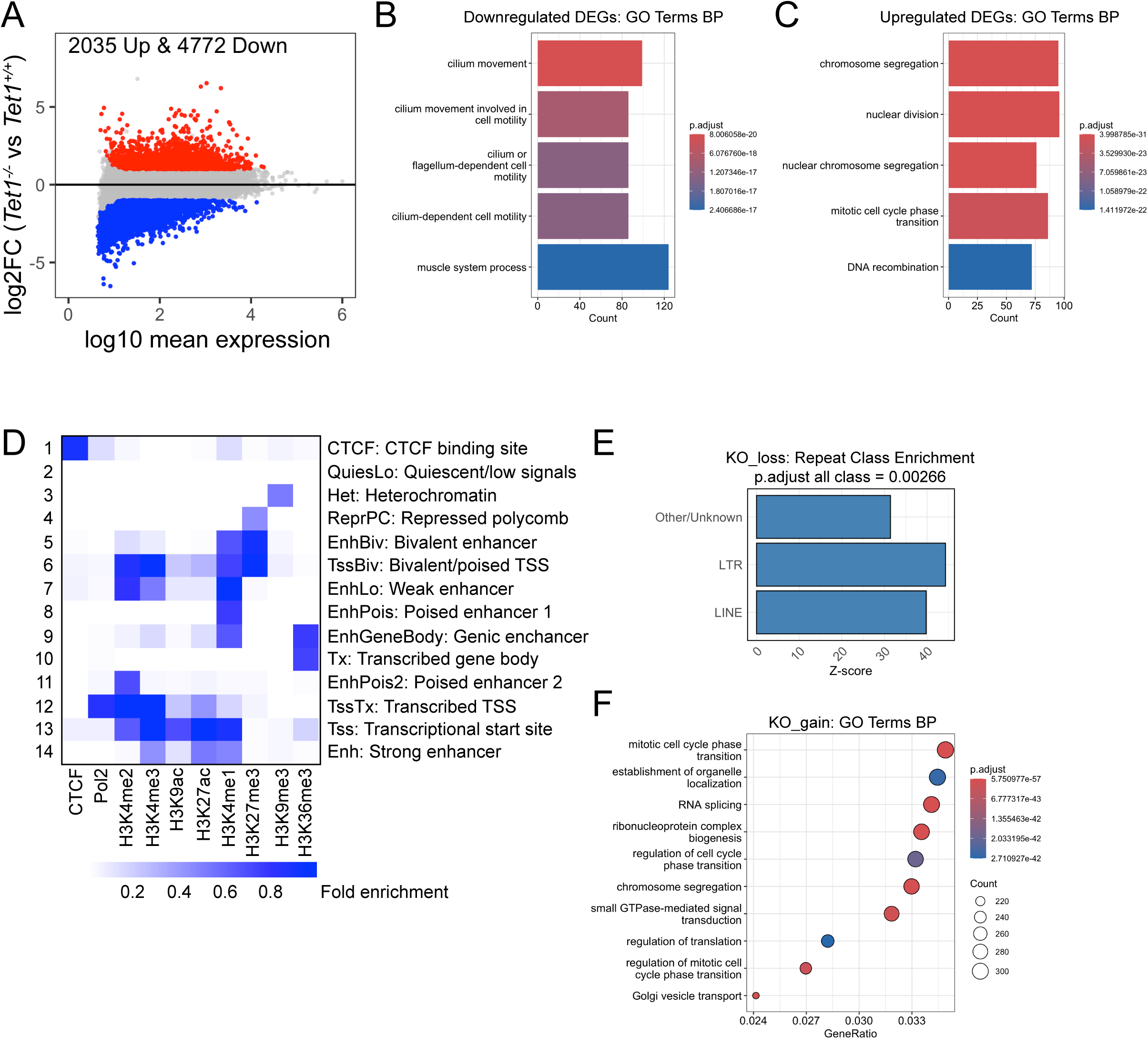
Investigation into the potential role of H3K4me3 changes in *Tet1^−/−^* prospermatogonia. A) MA-plot for differential expression analysis of total RNA-seq in *Tet1^−/−^* prospermatogonia vs WT (n=3, FDR ≤ 0.05, ≥2-fold change). B) Top five enriched GO biological process terms for downregulated DEGs in *Tet1^−/−^* prospermatogonia. C) Top five enriched GO biological process terms for upregulated DEGs in *Tet1^−/−^* prospermatogonia. D) Chromatin states assigned by ChromHMM using public datasets for CTCF, Pol II, and histone modifications across spermatogonial stem cells (SSC), spermatogonia (SPG), pachytene spermatocytes (PS), round spermatids (RS), and sperm (datasets used are in supplemental table). Chromatin state assignment is based on Encode consortium convention (Ernst and Kellis 2017). E) Classes of repetitive elements enriched in regions with H3K4me3 loss. F) Top ten enriched GO biological process terms for regions with H3K4me3 gain.

**Supplemental Figure 4.**
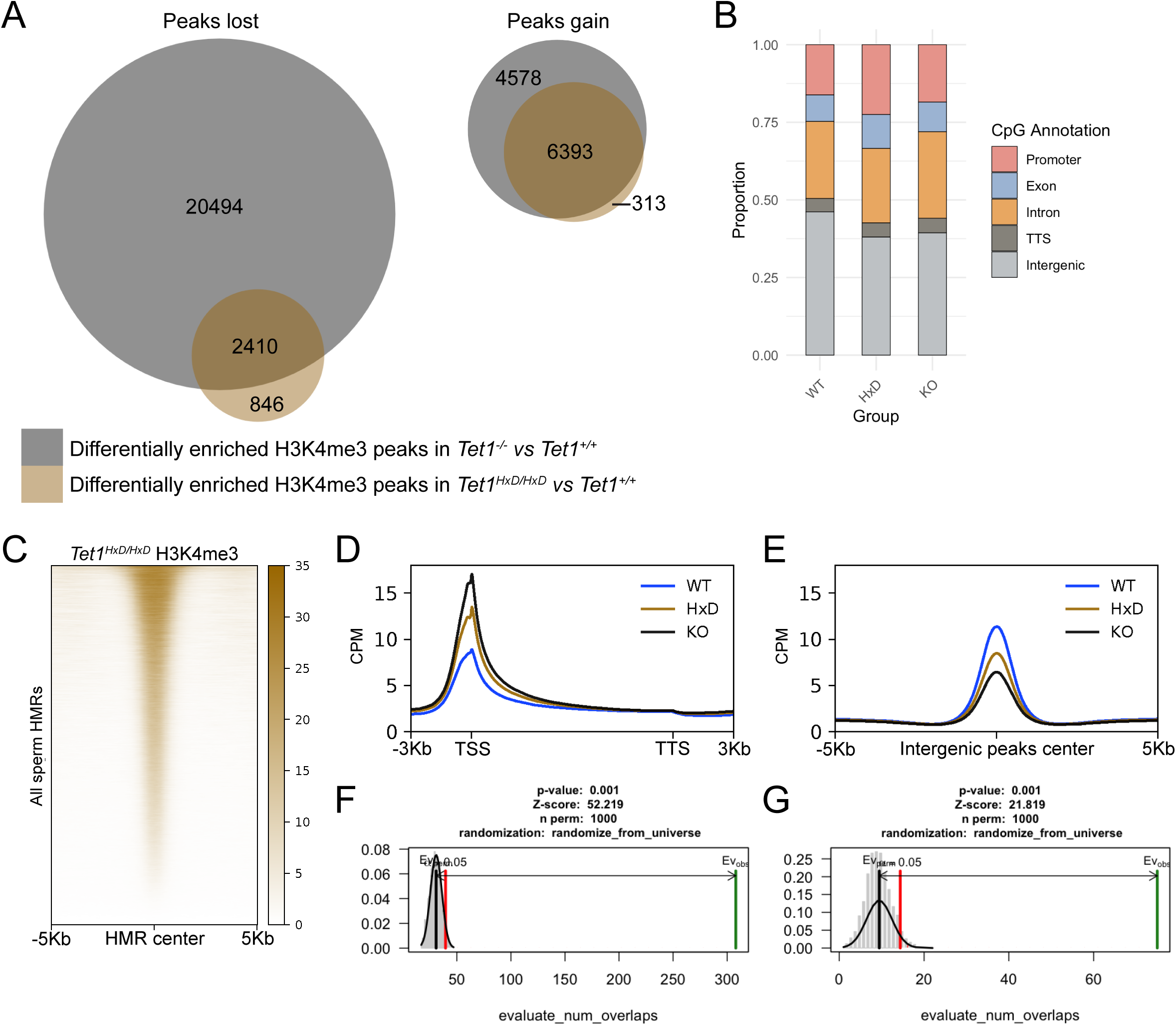
Extended CUT&RUN analysis for H3K4me3 in *Tet1^HxD/HxD^* prospermatogonia. A) Venn diagrams of differentially enriched peaks in *Tet1^−/−^* vs *Tet1^+/+^* and *Tet1^HxD/HxD^* vs *Tet1^+/+^* prospermatogonia (DiffBind, FDR ≤ 0.05, ≥1.5-fold change). B) Genomic compartmentalization of consensus peaks (≥2 of 3 replicates) in *Tet1^+/+^*, *Tet1^HxD/HxD^*, and *Tet1^−/−^*prospermatogonia. C) Heatmaps of H3K4me3 signals in E17.5 *Tet1^HxD/HxD^* prospermatogonia overlapping sperm HMRs defined by WGBS (GSE56697; Wang et al. 2014). Plots represent averages of 3 replicates. D–E) Metaplots of H3K4me3 signals over genic regions (D) or intergenic consensus peaks (E) for *Tet1^+/+^*, *Tet1^HxD/HxD^*, and *Tet1^−/−^*prospermatogonia. F-G) The significance of enrichment of TET1^HxD^-rescued (F) or TET1^HxD^-not rescued (G) DMRs within regions where H3K4me3 was restored to WT levels in *Tet1^HxD/HxD^*prospermatogonia following observed depletion in *Tet1^−/−^* prospermatogonia. In the graph, the grey-shaded region under the curve represents the distribution of the number of overlaps between 1000 randomized DMRs (within the array coverage) and the fixed set of H3K4me3-restored regions, values above the red line correspond to the area with a significance of p < 0.05. The solid green line represents the observed number of overlaps between rescued or not-rescued DMRs and H3K4me3-restored regions. Z-scores represent the strength or standardized magnitude of enrichment of overlap.

## Notes

### Competing Interest Statement

The authors have declared no competing interest.

### Summary of Updates

This paper has been update following peer-review and submitted as final version prior to journal edits.

